# IP_3_R1 is required for meiotic progression and embryonic development by regulating mitochondrial calcium and oxidative damage

**DOI:** 10.1101/2024.05.20.594928

**Authors:** Chang Zhang, Xiaoqing Sun, Deyi Wu, Guoxia Wang, Hainan Lan, Xin Zheng, Suo Li

**Author notes:** **Correspondence:** Suo Li. These authors contributed equally to this work (Chang Zhang, Xiaoqing Sun).

## Abstract

Calcium ions (Ca^2+^) regulate cell proliferation and differentiation and participate in various physiological activities of cells. The calcium transfer protein inositol 1,4,5-triphosphate receptor (IP3R), located between the endoplasmic reticulum (ER) and mitochondria, plays an important role in regulating Ca^2+^ levels. However, the mechanism by which IP_3_R1 affects porcine meiotic progression and embryonic development remains unclear. We established a model in porcine oocytes using siRNA-mediated knockdown of IP_3_R1 to investigate the effects of IP_3_R1 on porcine oocyte meiotic progression and embryonic development. The results indicated that a decrease in IP_3_R1 expression significantly enhanced the interaction between the ER and mitochondria. Additionally, the interaction between the ER and the mitochondrial Ca^2+^ ([Ca^2+^]_m_) transport network protein IP_3_R1-GRP75-VDAC1 was disrupted. PLA decreased IP_3_R1, weakened the pairwise interaction between IP_3_R1-GRP75 and VDAC1 and significantly enhanced the interaction between GRP75 and VDAC1, resulting in the accumulation of large amounts of [Ca^2+^]_m_. These changes led to mitochondrial oxidative stress and reduced ATP production, which hindered the maturation and late development of porcine oocytes and caused apoptosis.

## INTRODUCTION

Cell proliferation is an important characteristic of cell life activities and is the basis of biological growth, development and reproduction. The smooth differentiation and appreciation of oocyte and embryo development determine an individual’s life. As the basis of embryonic development, oocytes provide large amounts of essential nutrients. Therefore, the developmental potential of oocytes is directly related to the subsequent development ability of fertilization and embryos, directly affects the fertility of females, and even determines the fate of newborn individuals. However, good oocyte development requires a balance among various energy metabolism pathways. In addition to the need for an external nutrient supply, the maintenance of homeostasis in the intracellular environment is also crucial.

Calcium homeostasis is essential for maintaining cellular homeostasis and maintaining intercellular function^[1, 2]^.Due to the limited acquisition route and limited number of mature oocytes in vivo, in vitro maturation (IVM) culture has become another important way to obtain mature oocytes. However, compared with those in the body, oocyte IVM culture from the body follicle microenvironment is highly vulnerable to the influence of external environmental changes. When cells are stimulated by an imbalance in intracellular calcium homeostasis, a series of cascade reactions and organelle network disorders occur, leading to apoptosis, which affects oocyte development. During the in vitro maturation of porcine oocytes, mitochondrial damage and endoplasmic reticulum stress are accompanied by changes in the cytoplasmic calcium ions of oocytes, resulting in low-quality and low-efficiency porcine oocyte in vitro maturation, thereby affecting the developmental potential of oocytes.

The endoplasmic reticulum (ER) is crucial in cells^[3]^ and is involved in calcium storage, protein synthesis, and lipid metabolism^[4]^. The ER is a significant site of Ca^2+^ storage, and mitochondria are the main effectors of Ca^2+^ uptake and apoptosis^[5]^. Normally, intracellular Ca^2+^ homeostasis depends primarily on three calcium channels within the lumen of the endoplasmic reticulum: RyR^[6]^, IP_3_R, and SERCA. In general, the RyR^[6]^ and IP_3_R receptors are responsible for the release of Ca^2+^ from the ER lumen into the cytoplasm, while the SERCA calcium pump takes up cytoplasmic Ca^2+^ into the ER ^[4]^. This process continues uninterrupted, maintaining the dynamic equilibrium of Ca^2+[7]^ directly into the mitochondria through the cytoplasm, indicating the presence of Ca^2+^ transporters between the endoplasmic reticulum and mitochondria that regulate the internal communication of calcium ions^[8]^. Hiroki Akizawa *et al*. also showed that IP_3_R1 is closely related to calcium oscillation, affecting the release of calcium ions in the endoplasmic reticulum in mice^[9]^.

Different organelles coordinate with each other to maintain cellular physiological activities, including oocyte physiological activities. Organelles can be generated through physical connections between organelle membranes^[10]^. The most typical physical connection between organelles is the formation of a mitochondria-associated membrane (MAM) between the outer mitochondrial membrane and the endoplasmic reticulum membrane^[11, 12]^.

The proteins involved in Ca^2+^ transport in MAMs include proteins associated with the endoplasmic reticulum^[3]^, the outer mitochondrial membrane, and the inner mitochondrial membrane. The calspirin inositol 1,4,5-triphosphate (IP_3_R) receptor in the endoplasmic reticulum is a component that regulates homeostasis between the endoplasmic reticulum and mitochondrial Ca^2+[13]^.

IP_3_R1 is the most highly expressed subtype in oocytes and is distributed across the endoplasmic reticulum membrane^[14]^. During oocyte maturation, the endoplasmic reticulum is reorganized and divided into clusters, and IP_3_R1 is transferred from the blastocyst region to the adjacent agglutinated plasma membrane region^[15]^. This unique IP_3_R1 cortical cluster and complete recombination of the ER in IBD oocytes are necessary for repeated calcium waves during fertilization^[16]^. In addition, in an investigation in mice, IP_3_R1 expression was shown to improve mitochondrial function and alleviate oocyte damage in obesity^[17]^. These results further suggest that ER IP_3_R1 plays a significant role in the regulation of Ca^2+^ homeostasis and the function of the ER and mitochondria^[18]^.

Our previous study revealed that IP_3_R deletion significantly affects in vitro maturation^[2]^, but the specific mechanism underlying the effect of IP_3_R on porcine oocytes is unknown. In particular, the interaction between mitochondria and the endoplasmic reticulum in oocytes, the role of the calcium transport interactor IP_3_R1 on MAMs and the interaction network of IP_3_R1-GRP75-VDAC1 are not known. This study analysed the role of IP_3_R1 in the regulation of calcium homeostasis in porcine oocytes and revealed the interaction of mitochondria with the ER and the participation of the IP_3_R1-GRP75-VDAC complex in oocyte development.

## RESULTS

### IP_3_R1 interference arrests meiotic progression in porcine oocytes

To study the effect of IP_3_R1 on porcine oocytes, we silenced IP_3_R1 with siRNA and used NC-siRNA as a control transfection reagent. To determine the transfection efficiency, after transfection with fluorescently labelled siRNA, the oocytes were collected at 12 h and assessed under a fluorescence microscope. As shown in the figure, the transfected oocytes exhibited green fluorescence, confirming that the transfection was effective. IP_3_R1 expression was measured by qPCR, which showed significantly reduced mRNA expression levels of IP3R after interference, which could be used in subsequent experiments (Figure 1A-B).

**FIGURE 1.**
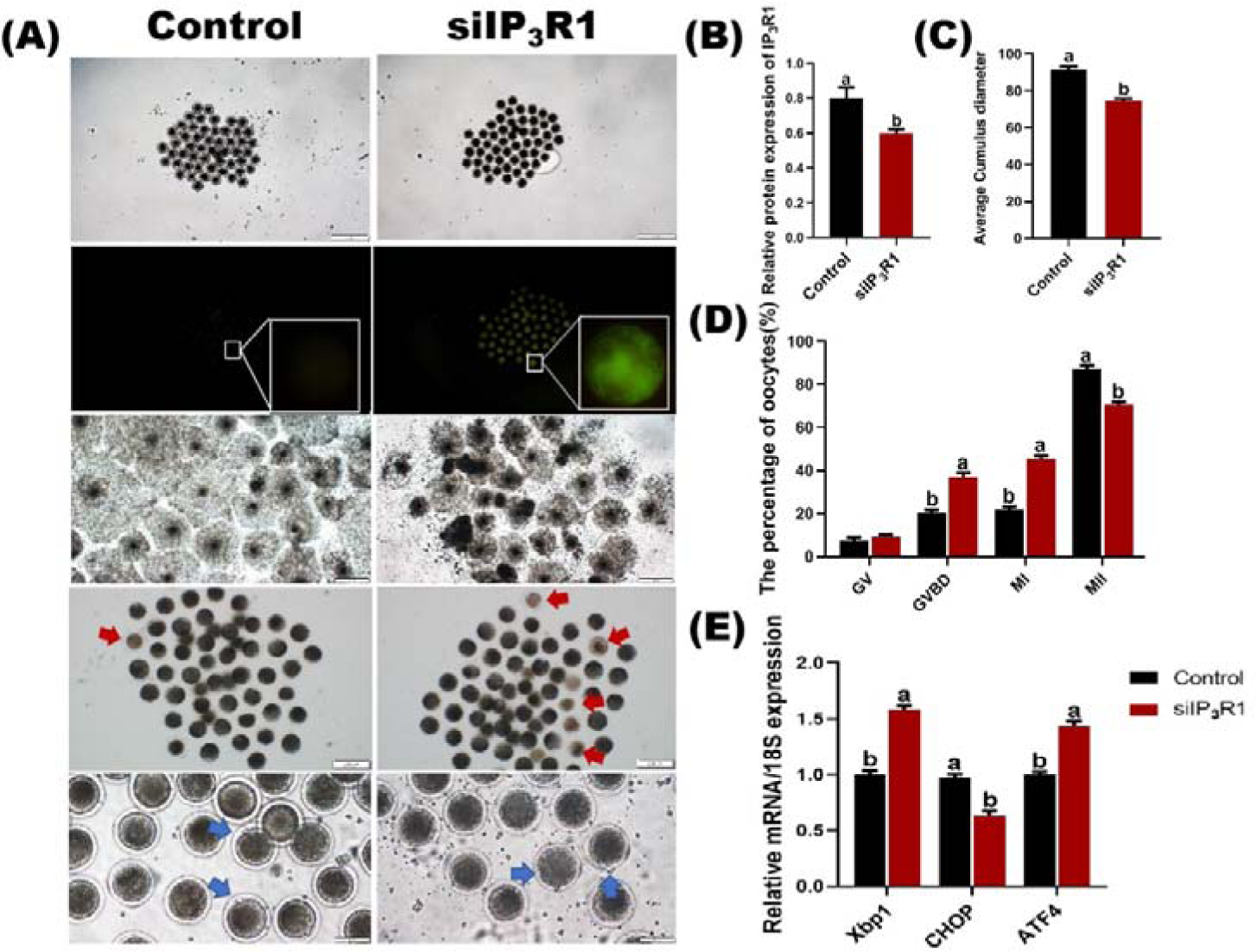
Effect of siIP_3_R1 on porcine oocyte maturation. (A) Transfection efficiency was assessed using NC-siRNA as a control. After transfection, the oocytes were collected 12 h later, and the transfection efficiency was assessed via fluorescence microscopy. Green: transfected fluorescently labelled siRNAs; oocyte cumulus diffusion and polar body discharge after IP_3_R1 interference. Red arrows indicate dead cells, and blue arrows indicate polar bodies expelled by mature cells. (B) siIP_3_R1 interference effect on mRNA detection. (C) First polar body expulsion and mortality after IP_3_R1 interference. Bar =100 μm. (D) Oocyte progression at various stages of meiosis. (E) The expression of ER-related genes after IP_3_R1 interference induced the UPR.

Subsequently, we further examined the effect of IP_3_R1 interference on oocyte maturation. siIP_3_R1 was added to porcine oocyte culture medium, and the maturity of the oocytes was assessed. After 44 h of IVM culture, the cumulus cells in the control group exhibited greater expansion, while the cumulus diffusion diameter in the siIP_3_R1 treatment group was significantly lower, and the diffusion was uneven (Figure 1A-C). After removing cumulus cells, the number of apoptotic cells in the siIP_3_R1 group was much greater than that in the control group. At the same time, mature cells in the siIP_3_R1 group had small, flattened polar bodies, and mature cells in the control group had large, rounded polar bodies. (Figure 1A, D).

The progression of oocyte meiosis was subsequently observed, and the number of oocytes in the siIP_3_R1 group was greater than that in the control group at the GV, GVBD and MI stages, but the number of mature oocytes in the siIP_3_R1 group was significantly lower than that in the control group at the MII stage.

Since IP_3_R1 is distributed on the ER membrane, we detected the UPR-associated genes Xbp1, Chop, and ATF4. Xbp1 and ATF4 gene expression was significantly increased, and Chop gene expression was significantly decreased (*P<0.05*) after IP_3_R1 interference (Figure 1E), indicating that IP_3_R1 interference caused ERS in oocytes.

### IP_3_R1 interference promotes mitochondrial and endoplasmic reticulum interactions

Since IP_3_R1 interference has previously been shown to cause an ERS response and mitochondrial oxidative damage, the interaction between the ER and mitochondria was further examined. Transmission electron microscopy (TEM) was used to measure the change in distance between these two organelles. Transmission electron microscopy (Figure 2A) revealed that the distance between the ER and mitochondria was significantly reduced after siIP_3_R1 treatment (*P<0.05*).

**FIGURE 2.**
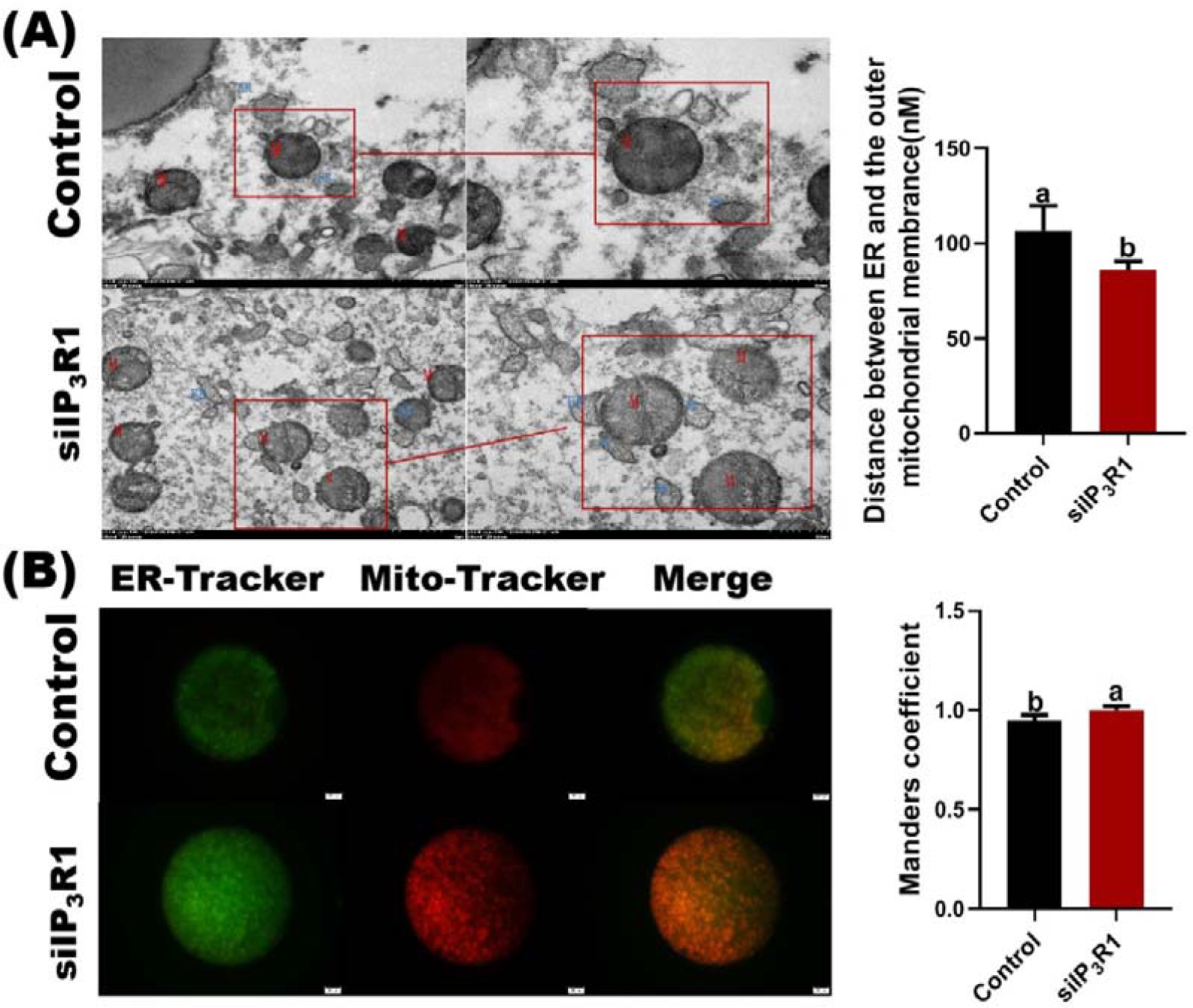
Effect of IP_3_R1 interference on the distance between mitochondria and the endoplasmic reticulum. (A). Oocytes (red indicates mitochondria; blue indicates the endoplasmic reticulum). Scale bar, top = 1 μm, middle = 2 μm, bottom = 500 nm. Measurement of endoplasmic reticulum-mitochondrial interactions. The data are presented as the mean ± standard error of the mean; n=4 independent experiments; *P<0.05 and #P<0.01; and analysis of variance was performed by Bonferroni post hoc correction. Measurement of endoplasmic reticulum and mitochondrial morphology and interactions. Green: endoplasmic reticulum. Red: Mitochondria. The Mandel coefficient indicates the strength of the interaction between two variables. Bar = 500 μm.

Immunofluorescence detection revealed that the fluorescence signal between mitochondria and the ER was significantly enhanced after IP_3_R1 interference, and Mander’s coefficient was also significantly increased (*P<0.05*), indicating that IP_3_R1 interference significantly enhanced the interaction between the ER and mitochondria (Figure 2B). These results suggest that a decrease in the distance between the ER and mitochondria leads to enhanced organelle-to-organelle interactions.

### IP_3_R1 interference changes the interaction between the IP3R1, GRP75 and VDAC1 proteins

We conducted further studies on the calcium channel complex (IP_3_R1-GRP75-VDAC1) to explore whether IP_3_R1 plays a role in Ca^2+^ exchange between the ER and mitochondria. Immunofluorescence assays revealed that IP_3_R1, GRP75, and VDAC1 coexisted in siIP_3_R1-treated oocytes, suggesting that these proteins are good candidates for studying ER-mitochondria interactions (Figure 3A). Moreover, we detected the mRNA expression of related genes after IP_3_R1 interference and found that the mRNA levels of IP_3_R1 and GRP75 decreased significantly after interference, and the mRNA levels of VDAC1 also showed a decreasing trend (*P<0.05*) (Figure 3B).

**FIGURE 3.**
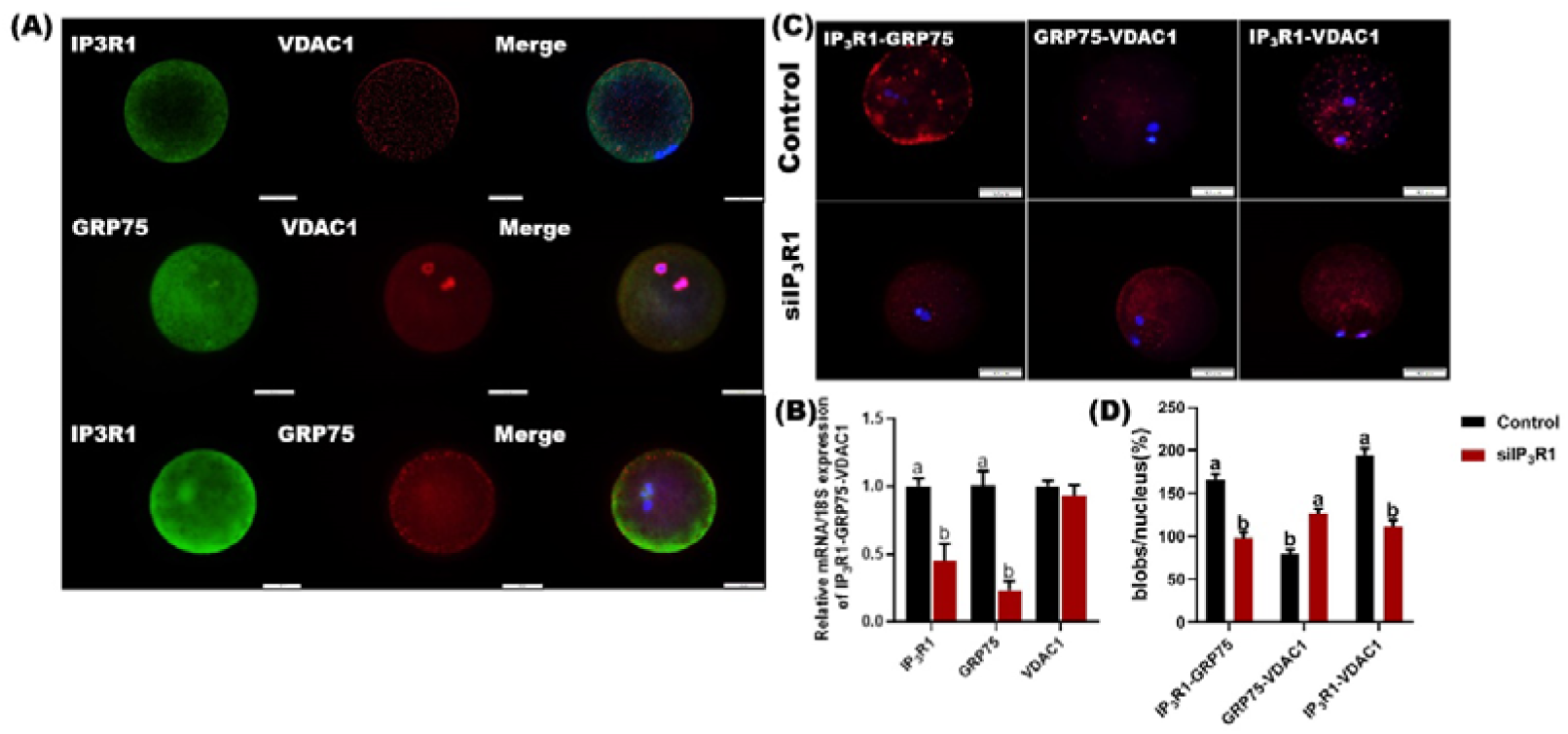
The IP_3_R1-GRP75-VDAC1 complex plays an important role in the interaction between the endoplasmic reticulum and mitochondria. **(A).** Conventional immunofluorescence confirmed the coexistence of IP_3_R1, GRP75 and VDAC1 in oocytes. Bar = 500 μm; (B). IP_3_R1-GRP75-VDAC1 mRNA expression. (C). In situ proximity ligation assays were used to monitor endoplasmic reticulum-mitochondrial interactions, which are associated with the genetic regulation of the IP_3_R1-GRP75-VDAC1 complex. The figure shows a typical PLA plot of the interaction between IP_3_R1-GRP75, GRP75-VDAC1, and IP_3_R1-VDAC1 after 44 h of oocyte culture with control or siIP_3_R1 treatment. Bar=500 μm. The values are presented as the mean ± standard error of the mean; n = 4 independent experiments; *P<0.05. PLA fluorescence and the distance between the endoplasmic reticulum and mitochondria were measured. The Manders coefficient and mean fluorescence intensity of the endoplasmic reticulum and mitochondria after siIP_3_R1 treatment.

Furthermore, we used the in situ proximity technique PLA to detect the protein interaction sites on this channel. Figure 3C shows a typical PLA plot of the interaction between IP_3_R1-GRP75, GRP75-VDAC1, and IP_3_R1-VDAC1 after 44 hours of oocyte culture in control or siIP_3_R1-treated oocytes. As shown in Figure 3D, after IP_3_R1 interference, the fluorescence signal between IP_3_R1-VDAC1 and IP_3_R1-GRP75 was significantly attenuated (*P<0.05*), and that between GRP75-VDAC1 was enhanced (*P*<0.05), indicating a decrease in protein‒protein interactions between GRP75-VDAC1 and a significant increase in protein‒protein interactions between IP_3_R1-VDAC1 and IP_3_R1-GRP75.

### IP_3_R1 interference causes abnormal Ca2+ transmission

The enhanced interaction between the ER and mitochondria can cause abnormal Ca^2+^ transmission, so we measured the effect of IP_3_R1 interference on [Ca^2+^]_i_, [Ca^2+^]_m_ and [Ca^2+^]_ER_ levels in porcine oocytes. As shown in FIGURE 4A-D, the siIP_3_R1 group had decreased [Ca^2+^]_i_ and [Ca^2+^]_ER_ compared to those of the control group. The [Ca^2+^]_m_ in the siIP_3_R1 group was significantly greater than that in the control group. The CALR mRNA level was also significantly increased (Figure 4E, *P<0.05*).

**FIGURE 4.**
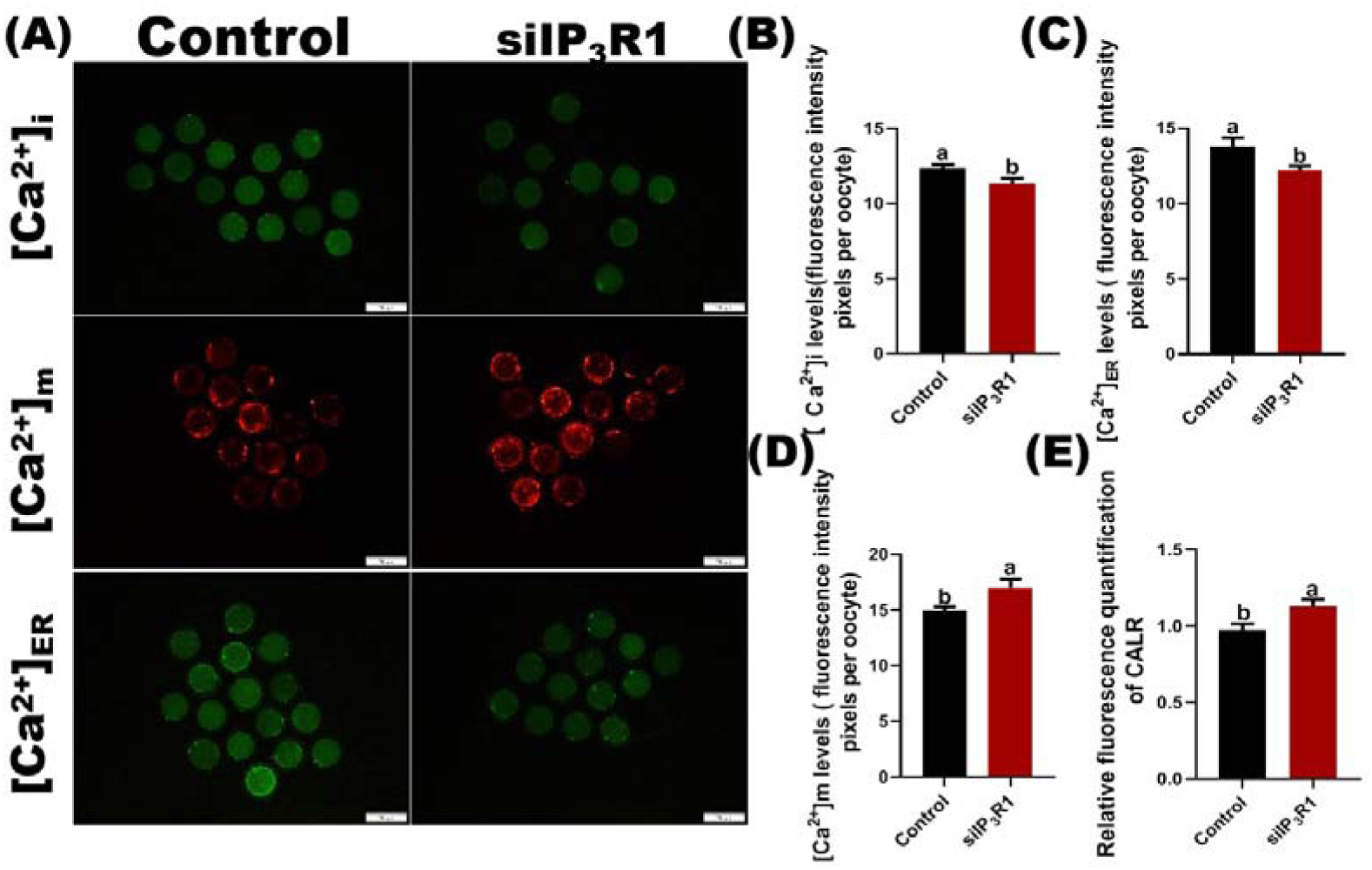
Effect of siIP_3_R1 on Ca^2+^ levels in porcine oocytes. **(A).** Fluorescence staining of [Ca^2+^]_i_, [Ca^2+^]_m_ and [Ca^2+^]_ER_. Bar =200 μm. (B). The fluorescence intensities of [Ca^2+^]_i_, [Ca^2+^]_m_ and [Ca^2+^]_ER_ after siIP_3_R1 treatment. Different superscripted letters denote a significant difference (a, b, P< 0.05). (C). The fluorescence intensities of [Ca^2+^] _ER_. (D). (E) The fluorescence intensities of [Ca^2+^]_m_. The fluorescence intensities of [Ca^2+^] _m_. (F). Superoxide dismutase (SOD) CALR mRNA levels were significantly increased.

### IP_3_R1 interference leads to mitochondrial oxidative damage and apoptosis

Exceptionally elevated [Ca^2+^]_m_ can cause mitochondrial dysfunction. Therefore, we examined the effects of IP_3_R1 interference on mitochondrial oxidative damage and apoptosis. As shown in Figure 5(A-B), oocytes in the control group had lower levels of ROS and higher levels of GSH, while oocytes inhibited by IP_3_R1 exhibited high levels of ROS and low levels of GSH signalling, suggesting oxidative stress following IP_3_R1 deficiency (control group *vs.* siIP_3_R1 group: ROS: 19.83 ±0.73 *vs.* 25.15±1.32; GSH: 56.43 ± 1.54 *vs.* 53.97 ±1.83; *P<0.05*).

**FIGURE 5.**
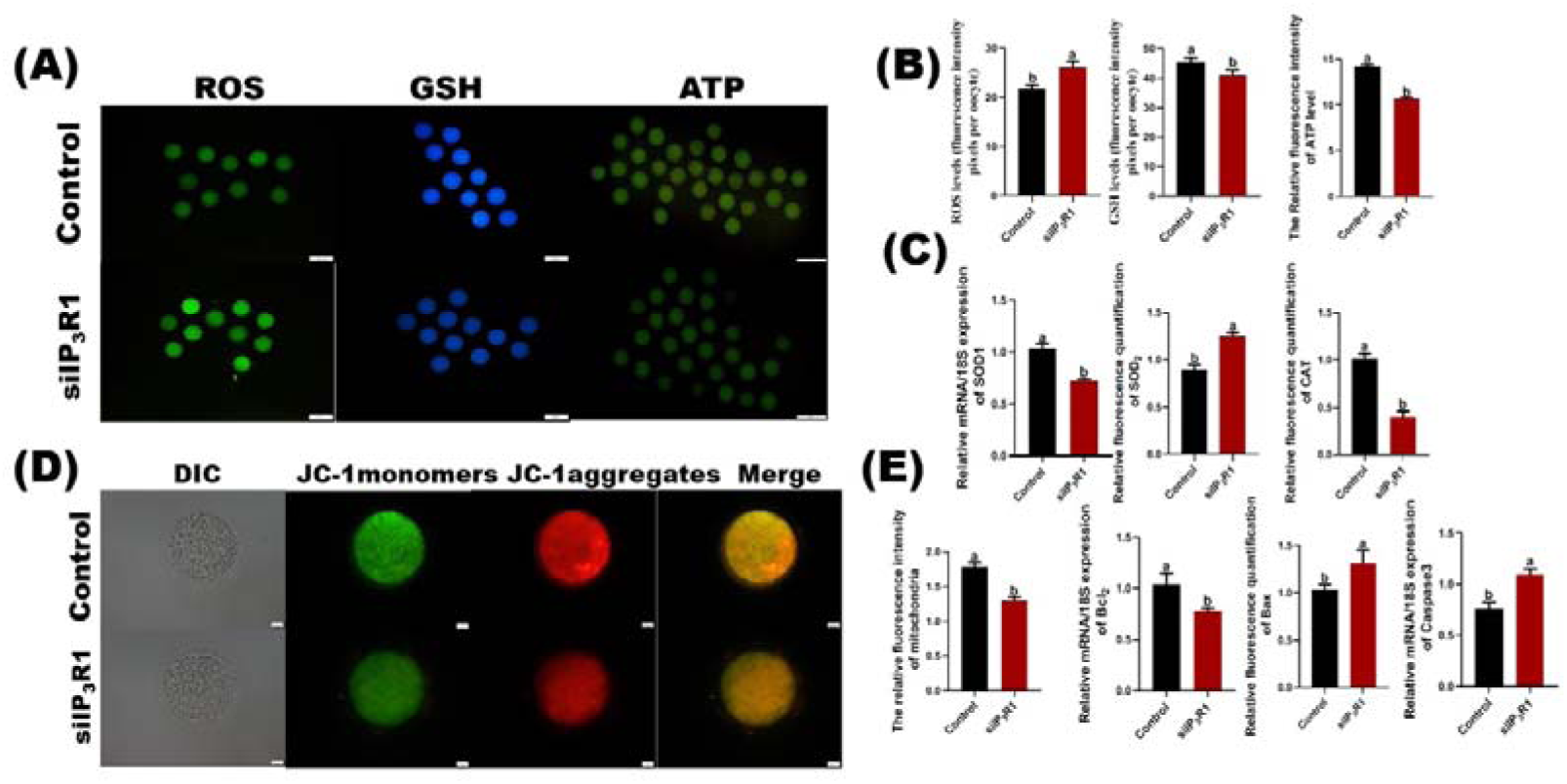
Effect of siIP_3_R1 treatment on mitochondria and apoptosis. (A). Fluorescence quantification of reactive oxygen species (ROS), glutathione (GSH) and ATP in each group after IP_3_R1 treatment (green, ROS). Blue, GSH. Bar = 100 μm; ROS and GSH fluorescence intensity analysis. The fluorescence intensity was much greater in the treatment group than in the control group. (B). The expression levels of ROS, GSH, and ATP mRNAs in each group. Bar = 100 μm. (C). The expression levels of the oxidative stress-related genes SOD1, SOD2 and CAT in each group. (D). Fluorescence quantification of the mitochondrial membrane potential (JC-1). (E). Genes associated with apoptosis and JC-1 mRNA expression.

In addition, the expression levels of SOD1, SOD2, CAT, Caspase3, Bax and Bcl_2_, which are related to oxidative stress and apoptosis, were detected. As shown in Figure 5(C, E), the expression levels of SOD1, CAT and Bcl2 decreased significantly, and the expression levels of SOD2, Caspase3 and Bax increased significantly in the siIP_3_R1 group. After siIP_3_R1 treatment, the fluorescence intensity of ATP was significantly weaker than that of the control group. The above results further confirmed that IP_3_R1 interference caused mitochondrial dysfunction damage, which caused oxidative stress and apoptosis, thereby affecting the IVM of porcine oocytes (Figure 5 A, D).

### IP_3_R1 is important for maintaining mitochondrial calcium levels

To further explore the effect of IP_3_R1 on [Ca^2+^]_m_ levels, the [Ca^2+^]_m_ chelator RR was used. Compared with those in the siIP_3_R1 group, the fluorescence intensity of [Ca^2+^]_m_ gradually returned to the level of that in the control group after the addition of RR, which was significantly lower than that in the siIP_3_R1 group, and [Ca^2+^]_ER_ and [Ca^2+^]_i_ were also tested. The results showed that the increase in [Ca^2+^]_ER_ and [Ca^2+^]_i_ in the RR group was significantly greater, and the levels of Ca^2+^ in the mitochondria, ER and cytoplasm returned to those in the control group, reaching the homeostatic equilibrium state of Ca^2+^ (Figure 6A-B). Further detection of oocyte maturation revealed that the diameter of the cumulus layer increased significantly, and the oocyte maturation rate and degeneration rate gradually returned to the levels of the control group (Figure 6C).

**FIGURE 6.**
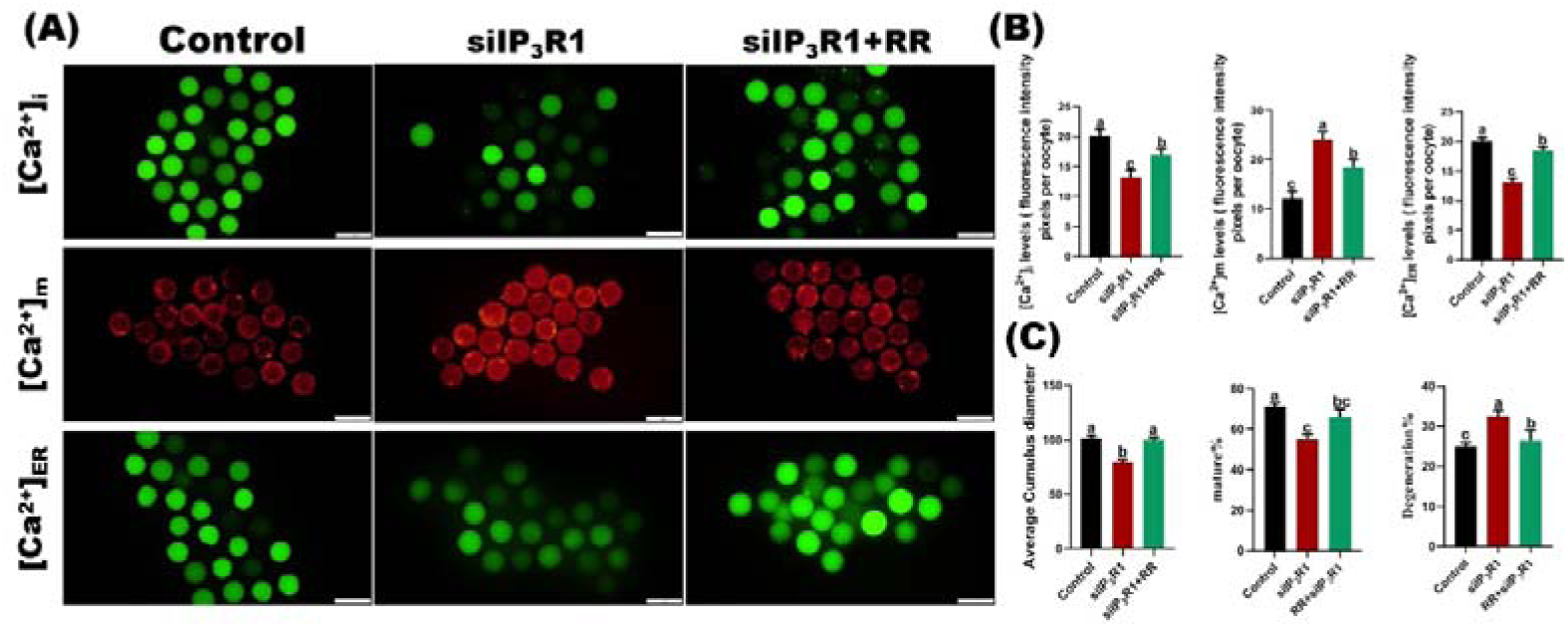
Effect of RR addition on calcium ion levels after siIP_3_R1 treatment. (A). Fluorescence staining of [Ca^2+^]_i_, [Ca^2+^]_m_ and [Ca^2+^]_ER_. Bar =200 μm. (B). The fluorescence intensities of [Ca^2+]^_i_, [Ca^2+^]_m_ and [Ca^2+^]_ER_ after RR addition. Different superscripted letters denote a significant difference (a, b, P< 0.05).

After [Ca^2+^]_m_ was confirmed to return to the level of the control group, mitochondrial function and ROS were further detected. After the addition of RR, the MitoTracker fluorescence signal in the siIP_3_R1 group was stronger, and the ROS fluorescence signal was significantly reduced (*P˂0.05*); that is, the mitochondrial level was significantly increased, and the ROS level was significantly reduced. This finding also demonstrated that the addition of RR after IP_3_R1 interference can help alleviate the mitochondrial damage and oxidative stress caused by IP_3_R1 interference (Figure 7).

**FIGURE 7.**
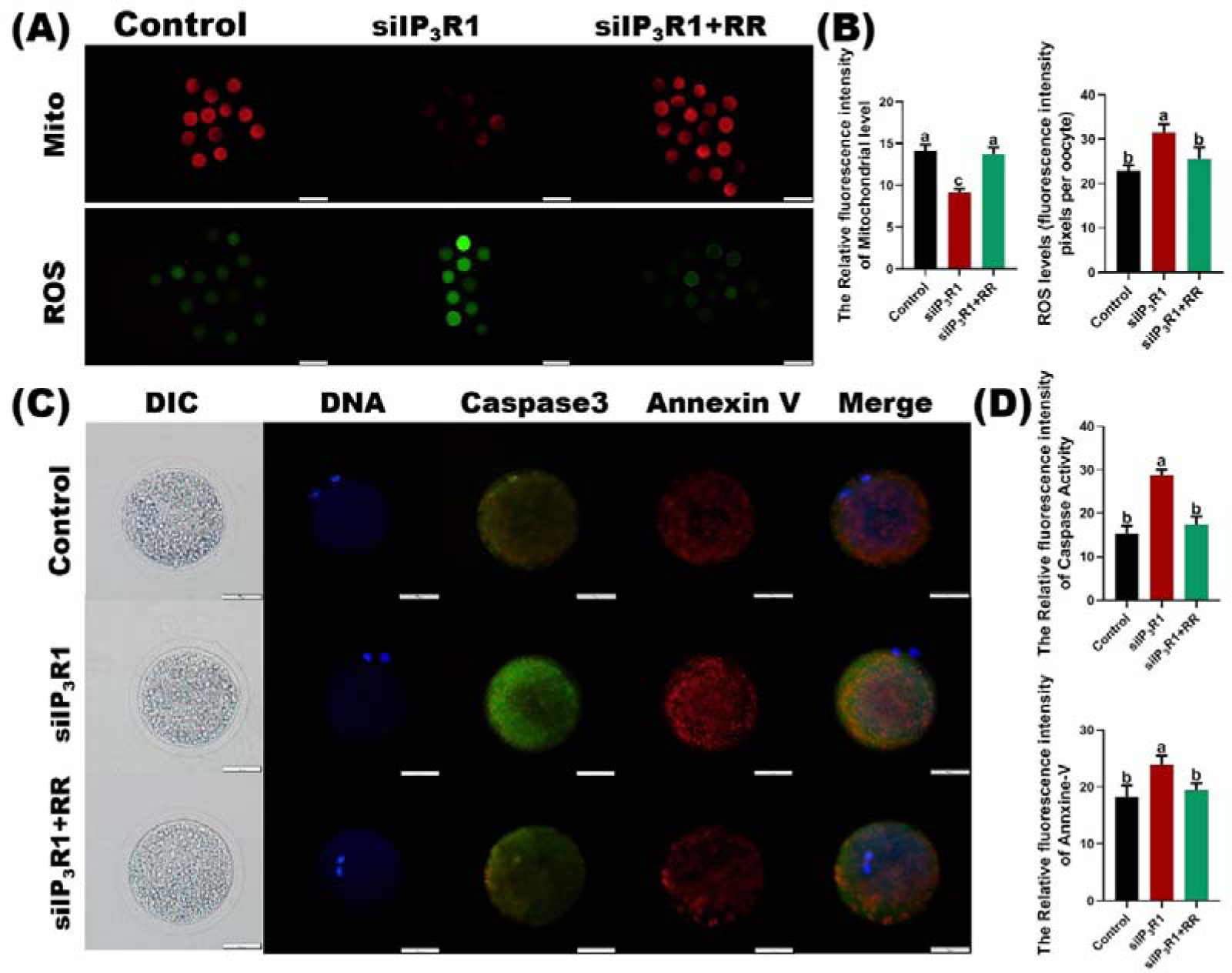
Effect of RR addition on mitochondria and apoptosis after siIP_3_R1 treatment. (A). Immunofluorescence staining of mitochondria and ROS after the addition of RR. Green: ROS. Red: Mitochondria. Bar = 100 μm. (B). The fluorescence intensities of mitochondria and ROS after RR addition. (C). Immunofluorescence staining for apoptosis after RR addition. Green: Caspase-3. Red: Annexin V. Bar = 500 μm. (D). The fluorescence intensities of Caspase-3 and Annexin V after RR addition.

Subsequently, the levels of Caspase3 and Annexin-V were detected, and we found that the fluorescence signals of the apoptotic mitochondrial pathway marker proteins Caspase3 and Annexin-V increased significantly after IP_3_R1 interference, and apoptosis was ameliorated after RR treatment. There was no significant difference between the RR group and the control group (Figure 7C-D, *P*<0.05).

### IP_3_R1 is important for maintaining mitochondrial oxidative damage

To further explore the effect of IP_3_R1 on mitochondrial oxidative stress, we utilized the ROS scavenger NAC. Compared with those in the siIP_3_R1 group, the immunofluorescence results showed that the mitochondrial fluorescence signal was more intense and the ROS fluorescence signal was less intense in the treatment group after the addition of NAC (Figure 8A, B), indicating that the addition of ROS scavengers after IP_3_R1 interference helped to attenuate the mitochondrial damage and oxidative stress caused by IP_3_R1 interference. Furthermore, the fluorescence signals of Caspase3 and Annexin-V increased significantly after IP_3_R1 interference, indicating an increase in apoptosis, and apoptosis was ameliorated after NAC treatment, gradually approaching the control group level (Figure 8C, D).

**FIGURE 8.**
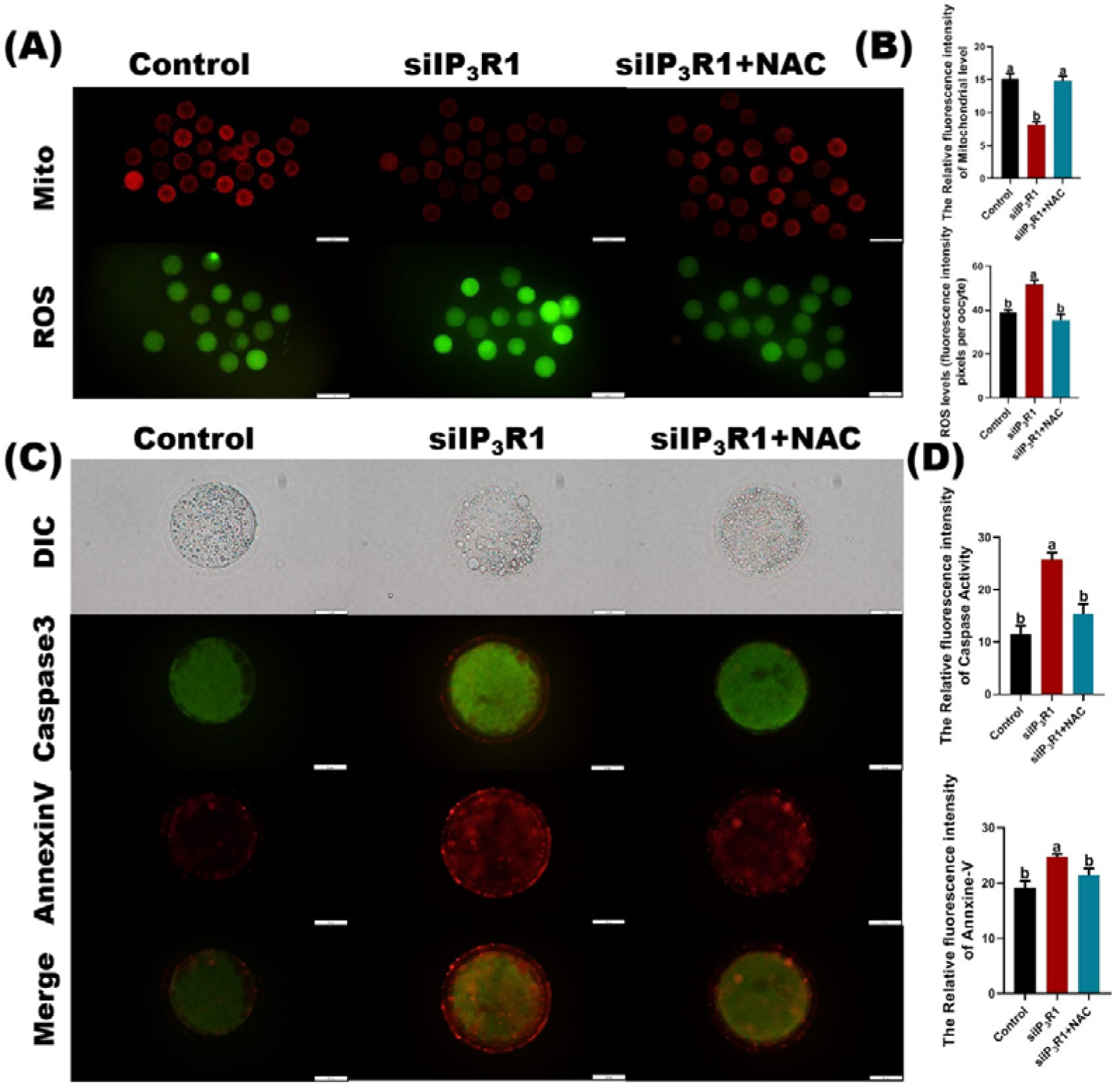
Effect of the ROS scavenger NAC on mitochondrial oxidative damage and apoptosis in oocytes. (A). The degree of cumulus cell expansion in each group improved after NAC supplementation, and the effect of NAC on mitochondrial function and ROS levels also improved. (B). Effect of NAC addition on apoptosis. Bar = 500 μm.

### IP_3_R1 interference leads to abnormal blastocyst development

To study the effect of IP_3_R1 interference on early embryonic development in pigs, PA was performed on mature oocytes, and cleavage and blastocyst morphology were recorded on days 2, 4, 5, and 7. We can see that the control group cells mostly begin to cleave the next day. In contrast, only a few cells in the siIP_3_R1 group began to undergo cleavage. By the fifth day, the control group had begun to divide and develop more blastocysts, while the siIP_3_R1 group had developed only a small number of smaller blastocysts. After continuing culture until the seventh day, most of the cells in the control group developed into blastocysts with a larger morphology, while the number of blastocysts in the siIP_3_R1 group was significantly less than that in the control group, the blastocyst morphology was smaller, and the blastocyst diameter was much smaller than that in the control group (control group *vs.* siIP_3_R1 group: 156.60 μm *vs*. 136.82 μm). According to the statistics, the cleavage rate (control group *vs.* siIP_3_R1 group: 88.87% *vs*. 71.46%) and blastocyst rate (control group *vs.* siIP_3_R1 group: 50.80% *vs*. 33.7%, (*P<0.05*)) in the siIP_3_R1 group were significantly reduced (Figure 9A, B).

**FIGURE 9.**
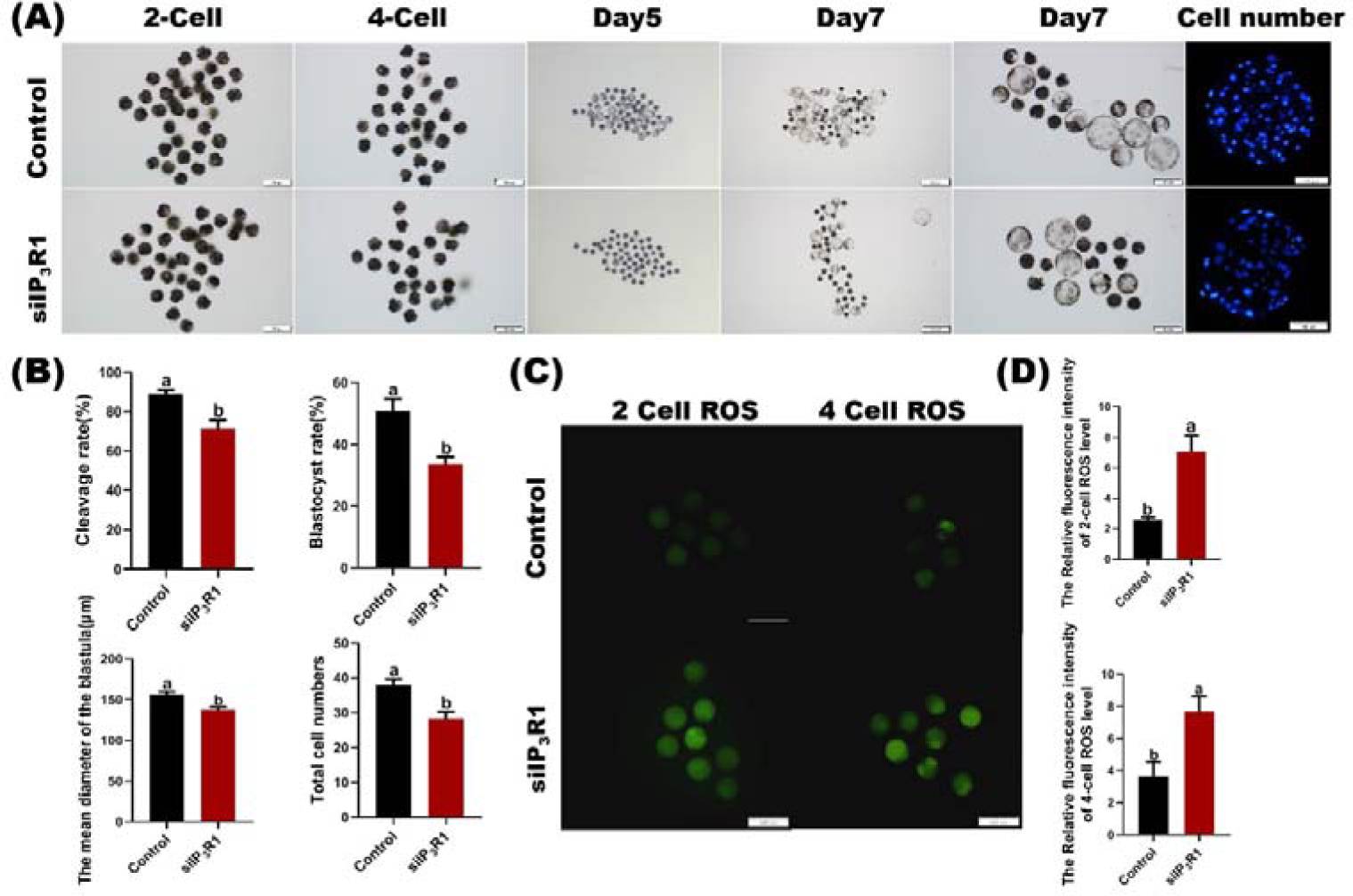
Effect of siIP_3_R1 treatment on embryonic development after PA treatment. (A). Cleavage and blastocyst developmental morphology in different groups at different stages. On days 5, 7, and 8, Hoechst 33342 staining (blue) was performed for the different treatment groups (100x magnification). (B). The ROS content, cleavage rate and blastocyst formation rate for the 2-cell and 4-cell phases.

Similarly, for the total number of cells (control group *vs.* siIP_3_R1 group: 38.03% *vs*. 28.76%), and based on the above results, we hypothesize that the addition of siIP_3_R1 affects the subsequent development of PA embryos.

To further investigate the effect of IP_3_R1 interference on the embryonic development of porcine oocytes, we performed ROS staining on cells subjected to parthenogenesis activation on days 2 and 4 after cleavage, and the ROS levels on days 2 and 4 were recorded and measured. The ROS fluorescence signal in the siIP_3_R1 group was significantly enhanced during the 2-cell and 4-cell stages, indicating increased oxidative stress (Figure 9C, D, *P<0.05*). This finding also demonstrated that the effect of IP_3_R1 on early embryonic development was related to mitochondrial oxidative damage.

## DISCUSSION

This study investigated the function of IP_3_R1 in porcine oocytes. We discovered that ERS was caused by the UPR unfolded protein response caused by IP_3_R1 interference, which subsequently enhanced the interaction between the ER and mitochondria via the ER-mediated IP_3_R1–GRP75–VDAC1 complex, resulting in [Ca^2+^]_m_ overload and mitochondrial oxidative damage, ultimately affecting the meiotic progression and embryonic development of porcine oocytes after PA.

Several studies have reported that mammalian oocytes almost abundantly express IP_3_R1^[19]^, and the most conspicuous mechanism regulating IP_3_R1 function is associated with oocyte maturation and the cell cycle. Ca^2+^ release through IP_3_R1 is greatly enhanced after the initiation of oocyte maturation and reaches a maximum during the meiotic M phase. Similarly, exit from M-phase and progression into interphase are associated with attenuation of Ca^2+^ responses^[20]^, which is accompanied by a pronounced loss of IP_3_R1 function. These findings indicate that IP_3_R1 plays a nonnegligible role in oocyte meiotic progression. Therefore, we transfected porcine oocytes with an IP_3_R1 plasmid to explore the effect of interference on the maturation and development of porcine oocytes and discovered that after IP_3_R1 interference, the oocyte diffusion diameter decreased, oocyte meiotic progression arrested in the GVBD or MⅠ stage, and MⅡstage oocytes significantly decreased. Our results and those of previous studies indicate an important role for IP_3_R1 in promoting meiotic progression in oocytes.

Moreover, interference causes the UPR to undergo an unfolded protein reaction, and interference with IP_3_R1 is closely related to the endoplasmic reticulum because disruption of ER homeostasis leads to the accumulation of unfolded or misfolded proteins. The endoplasmic reticulum is an important organelle for the IP_3_R-GRP75-VDAC1 calcium transport channel, and further studies of this channel complex were carried out. We discovered that the distance between the endoplasmic reticulum and mitochondria near this channel significantly decreased and that the distance between each mitochondrion increased due to IP_3_R1 interference. Our previous studies revealed that enhanced interactions between the endoplasmic reticulum and mitochondria affect calcium ion transport^[2]^. Impaired endoplasmic reticulum function results in calcium in the endoplasmic reticulum being released into the cytoplasm ^[8]^. After siIP_3_R1 transfection, the [Ca^2+^]_ER_ and [Ca^2+^]_i_ significantly decreased, and the [Ca^2+^]m significantly increased. [Ca^2+^]_ER_ may continue to be released outwards after endoplasmic reticulum dysfunction occurs, resulting in a surge of [Ca^2+^]_m_. Some similar studies have also demonstrated that in vitro culture away from the reproductive tract microenvironment causes insufficient nutrient supply to produce stress, leading to IP_3_ reduction, causing insufficient [Ca^2+^]_i_ oscillations, which is a potential cause of the failure of the oocyte-to-embryo transition^[21, 22]^. However, [Ca^2+^]_ER_ cannot be released when IP_3_R is inhibited by MAPK inhibitors during oocyte maturation ^[23]^.

Excess [Ca^2+^]m leads to increased apoptosis and impaired oocyte maturation in obese mice. However, the inhibition of IP_3_R1 may downregulate [Ca^2+^]_m_, improve mitochondrial function, and reduce apoptosis, thereby ameliorating obesity-induced oocyte damage and promoting in vitro maturation; these findings differ from the results of the present study, possibly because recent studies have focused on disrupting the role of IP_3_R in disease or lesional states ^[17] [24, 25]^. As a result, IP_3_Rs appear to function differently from those in normal cells. Arruda *et al.* ^[26]^ reported that in mice fed a high-fat diet, the increased expression of IP_3_R1 near the mitochondria results in high IP_3_R1 protein expression, resulting in the continuous pumping of calcium to mitochondria through the endoplasmic reticulum, ultimately leading to mitochondrial calcium overload, impaired mitochondrial function, and a disruption in metabolic homeostasis. Therefore, we hypothesize that downregulating IP_3_R1 expression leads to the aggregation and discretization of IP_3_R1, leading to endoplasmic reticulum stress, which also leads to the continuous pumping of calcium to mitochondria through the endoplasmic reticulum, ultimately leading to calcium overload. In this study, normal oocytes were used as the research object to analyse the differences between the normal state and the IP_3_R1 interference state and the key role of IP_3_R1.

The influx of Ca^2+^ into the mitochondrial matrix affects many aspects of mitochondria, such as impaired mitochondrial function, which affects mitochondrial ROS production^[27]^. A study by Hui Zhang *et al*. also showed that ROS accumulation caused DNA damage and apoptosis and that oxidative damage had a nonnegligible effect on oocyte development^[28]^. Mitochondrial dysfunction can also cause an associated response to oxidative stress. Therefore, we also tested the levels of ROS and GSH. The results showed that the level of ROS in oocytes after IP_3_R1 interference significantly increased and that the level of GSH significantly decreased; moreover, the expression of the oxidative stress-related genes SOD_1_, SOD_2_, Bcl_2_, Bax and CAT increased. Additionally, the ROS level in oocytes in the siIP_3_R1 group significantly increased, and the GSH level decreased to a certain extent. Furthermore, ATP expression significantly decreased in the siIP_3_R1 group. Notably, the results of some studies contradict the findings of the present study. In studies of mouse oocytes, [Ca^2+^]_m_ increased rapidly when [Ca^2+^]_ER_ was transferred to mitochondria via MAMs. Excess [Ca^2+^]m increases apoptosis and maturation retardation in obese mice. However, the inhibition of IP_3_R1 may downregulate [Ca^2+^]m, improve mitochondrial function, and reduce apoptosis. ^[17]^. The probable reason for these conflicting results is that recent research has focused on the role of IP_3_R1 ^[24, 25]^; therefore, its effects have been assessed in normal cells. However, Arruda *et al.*^[26]^ reported that in mice fed a high-fat diet, increased IP_3_R1 expression near mitochondria led to high expression of the IP_3_R1 protein, resulting in continuous pumping of calcium to mitochondria through the endoplasmic reticulum, ultimately leading to mitochondrial calcium overload, impaired mitochondrial function, and disruption of metabolic homeostasis. Therefore, we hypothesize that the downregulation of IP_3_R1 leads to the aggregation and discretization of IP_3_R1, leading to endoplasmic reticulum stress, which also leads to the continuous pumping of calcium to the mitochondria through the endoplasmic reticulum, ultimately leading to calcium overload. In this study, normal oocytes were used to compare differences between the normal state and the IP_3_R interference state, and the key role of IP_3_R was analysed.

We further studied the effect of IP_3_R1 interference on embryonic development. By observing cleavage and blastocyst morphology at different stages after parthenogenetic activation, we discovered that the cleavage rate in the interference group was significantly lower than that in the control group on days 2 and 4 and that the ROS level was also significantly increased, indicating that interference with IP_3_R1 had an effect on embryonic development. Further observation of blastocyst morphology on the 7th and 8th days showed that the control group had more blastocysts than did the interference group, which also demonstrated that interference with IP_3_R1 strongly affects the embryonic development of porcine oocytes.

In conclusion, our results showed that IP_3_R1 affects Ca^2+^ homeostasis in oocytes and induces [Ca^2+^]m overload and oxidative stress by regulating the Ca^2+^ transport network IP_3_R1-GRP75-VDAC1 between the ER and mitochondria, which promotes an increase in ROS levels and causes apoptosis, thereby affecting the meiotic progression and embryonic development of porcine oocytes (Figure 10).

**FIGURE 10.**
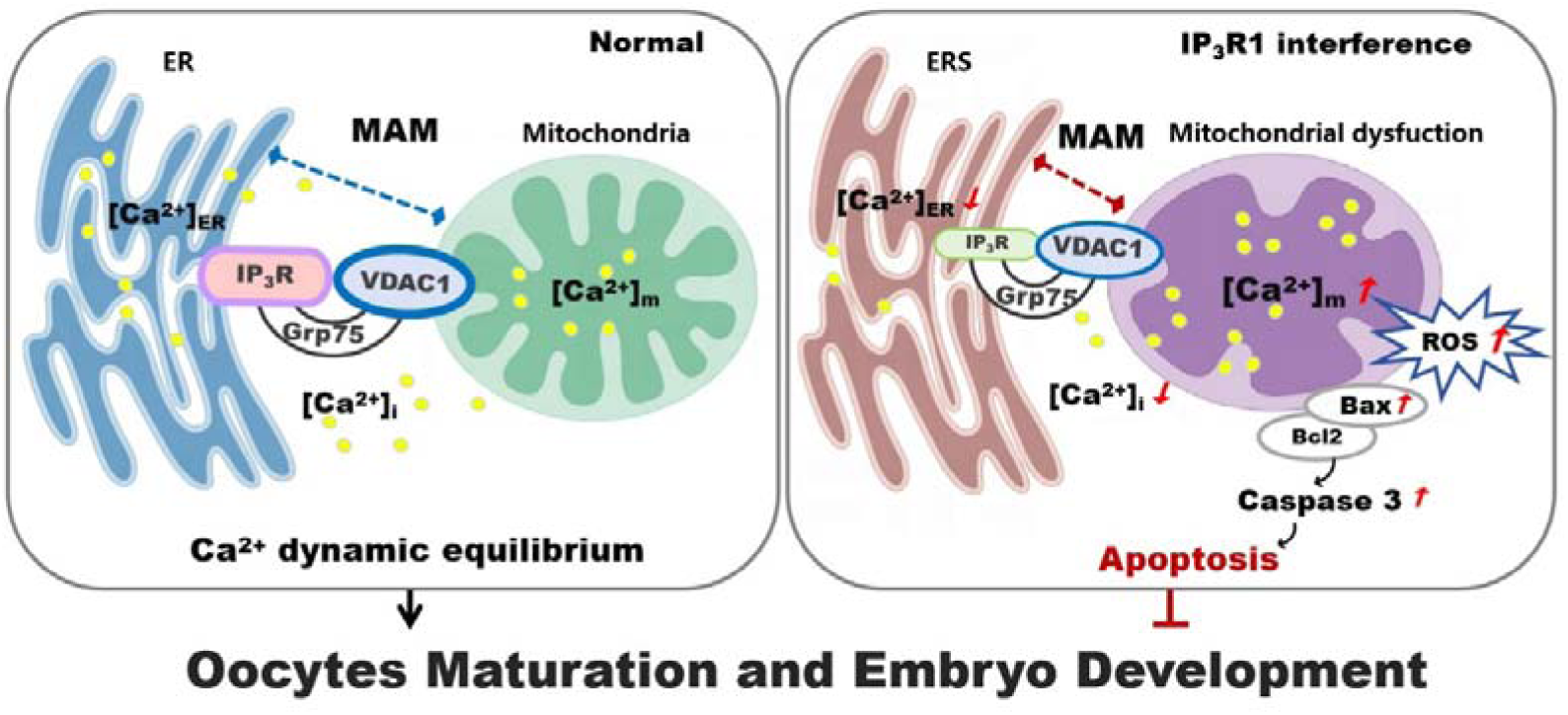
Graphical effect of IP3R1 on porcine oocyte maturation and embryonic development.

## CONCLUSION

IP_3_R1 regulates the dynamic balance of Ca^2+^ and the occurrence of oxidative damage by regulating the interaction between mitochondria and ER and the key IP_3_R1, GRP75 and VDAC1 factors, thus affecting the in vitro maturation of porcine oocytes and early embryonic development.

## METHODS AND MATERIALS

All animal experiments were conducted in accordance with the Guide for the Care and Use of Agricultural Animals in Research and Teaching. Unless otherwise specified, all the chemicals used were purchased from Sigma Chemical Co. (St. Louis, MO, USA).

### Oocyte harvest and in vitro maturation

The ovaries of prepubertal sows were collected at the local slaughterhouse, placed in an incubator containing 0.9% normal saline, and transported to the laboratory within 1 h. COCs were extracted from follicles (3-6 mm in diameter) using a 20 ml syringe. COCs were selected under a microscope. Serum gonadotropin (PMSG) (10 IU/ml) and human chorionic gonadotropin (HCG) (10 IU/ml) were added to TCM-199 and 10% pFF as IVM medium. COCs were cultured in a four-well culture dish at 38.5°C and 5% CO_2_ for 44 h to the MII stage. Cumulus cells were removed in 0.1% hyaluronidase, and the stripped oocytes were collected for subsequent analysis.

The detached oocytes were divided into different maturation stages (germinal vesicle (GV), germinal vesicle rupture (GVBD), metaphase 1 (MI) and metaphase II (MII)) for subsequent experiments.

### siIP_3_R1 primer construction and oocyte processing

siIP_3_R1 was designed by GenePharma (Suzhou, China). We used siIP_3_R1 to inhibit the activity of IP_3_R1 and investigated the effect of IP_3_R1 deficiency on porcine oocyte maturation. The sequence of the IP_3_R1 siRNAs used was as follows: siRNA-1: 50GCU CCA CAA UAA CCG GAA ATT - UUC CGG GUA UGG GAG CTT -30. We inhibited the activity of IP_3_R1 with siRNA to study the effect of IP_3_R1 interference on the in vitro maturation of porcine oocytes. siIP_3_R1 was mixed with the transfection agent RFectSP and dissolved in DMSO to 100 μL to preserve IP_3_R1, which was then diluted to 200 and added to 500 μL of IVM medium containing less than 0.2% DMSO.

### Mean diameter measurement of cumulus cell expansion

After oocyte maturation, the expansion of cumulus cells was observed under a microscope, and the two vertical diameters of COCs were measured with ImageJ software. A hallmark of maturation is the discharge of the first polar body (PB1), which is observed under the microscope.

### ROS and GSH level analysis

To mature naked eggs, ROS and GSH dyes were added, the samples were incubated in the dark for 30 minutes, and the stained samples were observed under a microscope.

### ATP staining

After removing the cumulus tissue from mature naked eggs, ATP rabbit monoclonal antibody and PBS+0.1% PVA were added at a ratio of 1:200 for mixing, the samples were added, the samples were incubated at 37°C for 15 min in the dark, and the images were observed under a microscope.

### Immunofluorescence staining

MII oocytes were collected. After the cumulus layer was removed, the fixative solution was added at room temperature for 30 minutes, and the samples were washed with PBS+0.1% PVA three times for five minutes each time. Subsequently, after permeabilization with 1% Triton X-100 for 10 min and blocking in blocking solution (PBS + 3% BSA) for 1 h, the oocytes were incubated with primary antibody overnight at 4°C [IP_3_R1: Invitrogen PAI-901/Grp75 rabbit (mouse) monoclonal antibody/VDAC1 rabbit (mouse) monoclonal antibody, 1:500]. After three washes with PBS + 0.1% PVA, the cells were incubated with a secondary antibody (goat anti-rabbit IgG A27034, goat anti-mouse Alexa Fluor 350, goat anti-rabbit Alexa Fluor 647, 1:250) for 1 h at room temperature in the dark, followed by Hoechst staining for 5 min. Laser scanning confocal microscopy was used to observe colocalization (Leica TCS SP5).

### Evaluation of mitochondrial and ER colocalization

Mature eggs were removed from the cell culture, stained with 0.5 μmol/L MitoTracker Red CMXRos (C1049B, Beyotime) and 0.5 μmol/L ER-Tracker Green (C1042S, Beyotime) and incubated at 37°C for 30 minutes in the dark. The interaction between mitochondria and the ER was evaluated by ImageJ software, and the colocalization of the mitochondrial ER was determined by Mande’s coefficient, as described in a previous report^[29]^.

### Mitochondrial membrane potential staining

Mature eggs were placed in the working solution of JC-1 for red fluorescence, mixed gently, and incubated at 38.5°C for 20 min in the dark. Subsequently, the slides were washed three times with 0.1% PVA. Hearst stain was added, and the glass slides were blocked, observed under a fluorescence microscope and photographed.

### Determination of [Ca^2+^]_i_, [Ca^2+^]_m_, and [Ca^2+^]_ER_

Pluronic F-127, Mag-Fluo-4 AM and Rhod-2 AM dyes were used to detect the [Ca^2+^]_i_, [Ca^2+^]_ER_, and [Ca^2+^]_m_, respectively. Mature naked eggs were washed three times with 0.1% PVA, placed in three calcium ion staining solutions, and incubated at 38.5°C for 15 min.

### Analysis of Caspase-3 and Annexin V contents

The supernatant was removed after centrifugation of the mature naked eggs, and the cells were gently resuspended in 194 μL of Annexin V-mCherry binding buffer. Then, 5 μL of Annexin V-mCherry and 1 μL of GreenNuc™ Caspase-3 Substrate were gently added, and the mixture was incubated for 30 minutes at room temperature in the dark. The cells were observed and imaged under a fluorescence microscope. GreenNuc-DNA™ is green fluorescence, and Annexin V-mCherry is red fluorescence.

### RNA isolation and quantitative real-time polymerase chain reaction

Forty mature oocytes were collected per group. RNA was extracted using an RNA kit (Invitrogen, Oslo, Norway). First-strand cDNA was synthesized using a cDNA synthesis kit (Plateau Biomedical Technology Co., Ltd., Dalian, China). Multiplex real-time polymerase chain reaction (RT\u2012qPCR) and qPCR were used to measure the levels of relevant δ. After the analysis of the melting curve, the gene expression level was analysed by the 2^-ΔΔ^ CT method. The expression levels of the target genes in each sample were normalized to those of 18S RNA (TABLE 1).

**TABLE 1.**
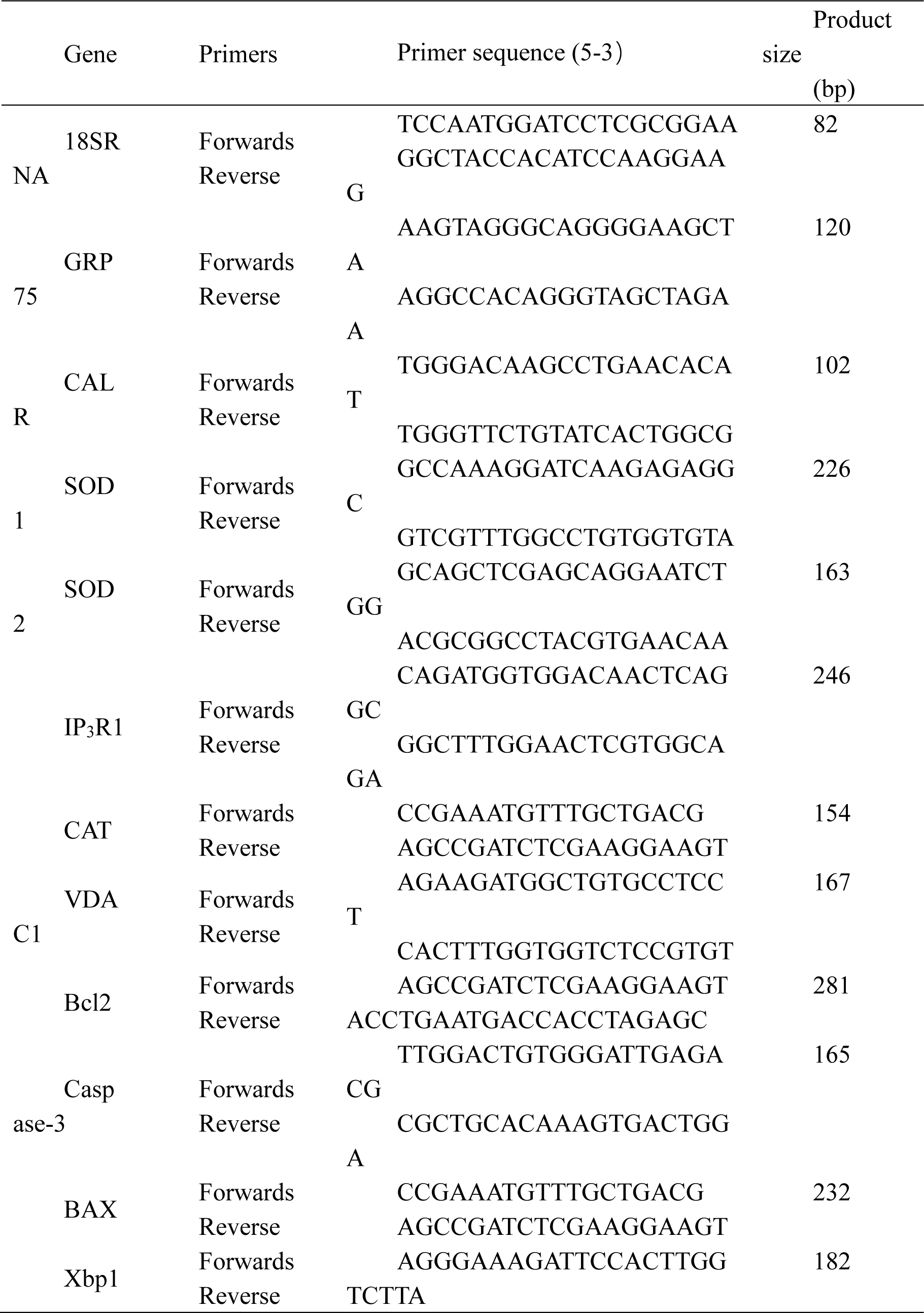
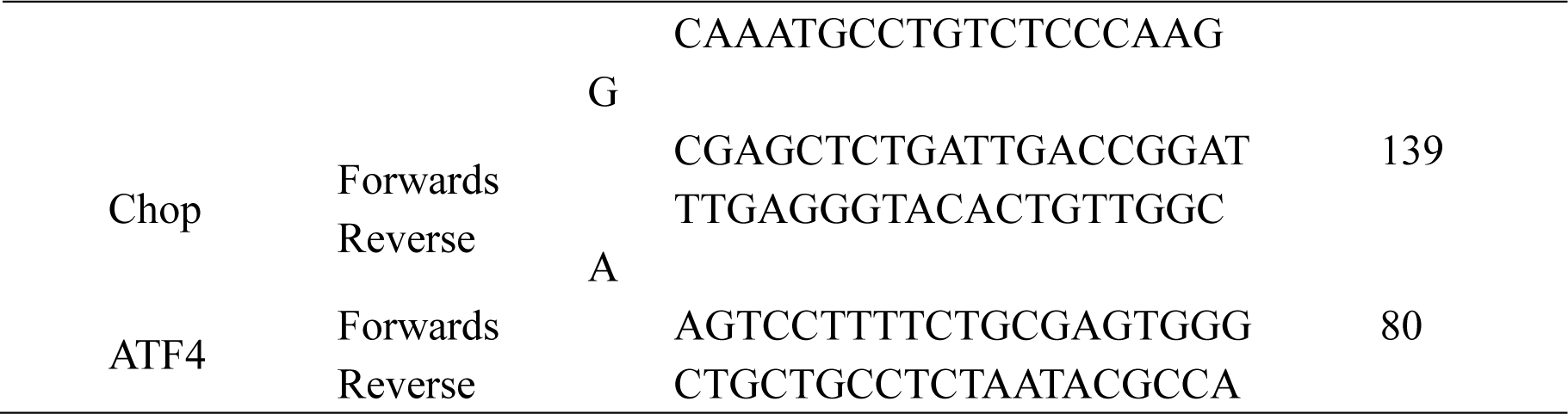
List of primers used for RT‒qPCR.

### Transmission electron microscopy

Approximately 120 mature oocytes of mung beans were collected, fixed with 2.5% glutaraldehyde and 1% permeable acid, dehydrated and incubated overnight in embedding medium. Sections were obtained with an ultramicrotome. After lead citrate staining, the sections were analysed under an electron microscope (10,000x).

### Proximity ligation in situ assay

The Duolink II in situ proximity ligation assay (PLA) (Olink Bioscience) enables microscopy to measure the proximity of two proteins and the potential interactions between them (<40 nm) as a single fluorescent spot. The mature oocytes were fixed at room temperature for 10 min, washed three times with PBS supplemented with 0.1% PVA after permeabilization for 30 min. The blocking solution was added to the sample, which was incubated at 37°C for 60 min, and the primary antibody (diluent) was added to the sample, which was subsequently incubated at 4°C in the dark overnight. Afterwards, the samples were washed 2 times with wash solution A for 5 min each, the probe was diluted 1:5 in buffer, the probe was added to the sample at 37°C, the probe was blocked for 60 min, the buffer A was washed twice for 5 min, the linked stock solution (Ligotion Buffer) was diluted 1:5 in high-purity water, the ligase was added to the ice box, the probe was blocked at 37°C for 30 min, the probe was washed twice with washing solution A for 5 min each time, the amplification solution was added to the ice box at 37°C, and the mixture was blocked for 60 min. The samples were washed twice with 1X buffer B for 10 minutes each and 0.01X buffer B for 1 minute, and the tablets were blocked with Duolink in situ mounting medium containing DAPI. Cells were imaged under an Olympus IX81 laser scanning microscope, and the signal was quantified using BlobFinder software and is expressed as the percentage of plaque in each nucleus compared to that in the control group.

### Parthenogenetic activation

Mature bare eggs were collected for IVM for later use. The collected oocytes were activated with a direct current pulse (1.2 kV/cm, 30 μs) utilizing a BXT Electro-Cell Manipulator 2001 (BXT Inc., San Diego, CA) in 0.28 mol/L mannitol containing 0.1 mM MgSO4 and 0.05 mM CaCl2. After PA, the oocytes were cultured in PZM-5 medium [4 mg/mL bovine serum albumin (BSA) and cytochalasin B (CB)] for 3 h and then transferred to in vitro culture medium in the same culture environment as IVM for 2 days to record their morphology and cleavage rate. The blastocyst formation rate was recorded on day 7.

### Statistical analysis

Images of mature oocytes after immunofluorescence staining were analysed with ImageJ. The mean values for the groups were analysed using GraphPad Prism 5. The data are presented as the mean ± standard error (SEM). Differences between two groups of data were analysed using Student’s t test with IBM-SPSS 23.0. *P* < 0.05 was considered to indicate statistical significance.

## ACKNOWLEDGEMENTS

This work was supported by the Jilin Scientific and Technological Development Program (No. YDZJ202301ZYTS330).

## AUTHOR CONTRIBUTIONS

LS and LHN designed the experiments; ZC and SXQ performed the majority of the experiments; WDY and WGX participated in the experiments; ZX contributed materials; SXQ analysed the data; and LS and ZC wrote the manuscript. All the authors approved the submission of the manuscript.

## DATA AVAILABILITY STATEMENT

The data that support the findings of this study are available from the corresponding author upon reasonable request.

